# Metabolic tagging reveals surface-associated lipoproteins in mycobacteria

**DOI:** 10.1101/2025.01.07.631728

**Authors:** Lia A. Parkin, Kindra L. Becker, Julian P. Maceren, Aseem Palande, Mary L. Previti, Jessica C. Seeliger

## Abstract

Mycobacteria such as the causative agent of tuberculosis, *Mycobacterium tuberculosis*, encode over 100 bioinformatically predicted lipoproteins. Despite the importance of these post-translationally modified proteins for mycobacterial survival, many remain experimentally unconfirmed. Here we characterized metabolic incorporation of diverse fatty acid analogues as a facile method of adding chemical groups that enable downstream applications such as detection, crosslinking and enrichment, of not only lipid-modified proteins, but also their protein interactors. Having shown that incorporation is an active process dependent on the lipoprotein biosynthesis pathway, we discovered that lipid-modified proteins are also located at the mycobacterial cell surface even though mycobacteria do not encode known lipoprotein transporters. These data have implications for uncovering a novel transport pathway and the roles of lipoproteins at the interface with the host environment. Our findings and the tools we developed will enable the further study of pathways related to lipoprotein function and metabolism in mycobacteria and other bacteria in which lipoproteins remain poorly understood.

## INTRODUCTION

Cell membranes are essential boundaries that maintain life. Accordingly, the inhibition of cell envelope biogenesis is a common mode of action among anti-bacterial drugs. Antibiotics against the human pathogen *Mycobacterium tuberculosis* (*Mtb*) are no exception, except that many target mycobacteria-specific pathways. In general, the mycobacterial cell envelope presents both opportunities and challenges in the fight against *Mtb* thanks to its unusual structure: In addition to a plasma membrane and peptidoglycan layer, mycobacteria are surrounded by an arabinogalactan layer that is covalently attached to ultra-long mycolic fatty acids that form the inner leaflet of an outer membrane, commonly called the mycomembrane due a diverse lipid composition that distinguishes it from the outer membrane of gram-negative bacteria. Given these distinguishing features, on the one hand, pathways unique to mycobacteria offer prospects for selective inhibition; on the other hand, the mycobacterial cell envelope limits the permeation of diverse small molecules into the cell^1–5^, including otherwise potent enzyme inhibitors, thus limiting cellular efficacy. The barrier role of the cell envelope has also been inferred from studies that have implicated lipid biosynthesis and metabolism genes in sensitizing mycobacteria to antibiotic activity^6–12^.

A strategy that incorporates both considerations is targeting processes in the cell envelope whose inhibition not only compromises bacterial survival, but also sensitizes mycobacteria to other antibiotics. However, exploration of this approach is constrained by our limited understanding of cell envelope biogenesis and cell envelope processes overall. Even our basic knowledge of what proteins reside within the cell envelope is limited by our incomplete understanding of export signals and by technical obstacles in accurately localizing proteins to the cell envelope, although we and others have developed methods that are helping to build a better picture of this subcellular proteome^13–16^. In contrast to these unknowns, lipid-modified proteins known as lipoproteins are well established cell envelope residents and the machinery that adds the post-translational modification is essential for *Mtb* survival^17–20^. In general, a conserved lipobox motif directs encoding proteins to the processing machinery^21,22^; while the lipobox is derived from studies in other bacteria, it is also found in mycobacteria. Based on bioinformatic analyses searching for this motif, lipoproteins make up ∼3% of the total *Mtb* proteome (>100 lipoproteins, as predicted in several reports^21,22^, including a recent review on mycobacterial lipoproteins^23^), consistent with the proportion predicted in other bacteria (1- 3%)^24^. In our recent report of proteins identified by subcellular compartment-specific protein tagging^13^, proteins encoding a lipobox motif constituted 19% of the detected cell wall proteome (49 of 254 proteins). Otherwise, only a handful of individual mycobacterial lipoproteins have been experimentally confirmed^23^.

*Mtb* lipoproteins are processed by a largely canonical pathway, based on bioinformatics and experimental validation in *Mtb* and other mycobacteria. In the consensus pathway, proteins with a lipobox motif are exported across the plasma membrane via primarily the Sec secretion system, although a minority may traverse the Tat system^25^. Integral membrane proteins further process the nascent lipoprotein in the periplasm (**Figure 1**). First, lipoprotein diacylglyceryl transferase (Lgt; Rv1614)^26^ catalyzes the addition of a diacylglycerol group via a thioether bond to the sidechain of a conserved cysteine (Cys) within the lipobox motif. Lipoprotein signal peptidase (LspA; Rv1539)^27^ cleaves upstream of the modified Cys. Apolipoprotein N-acyltransferase (Lnt; Rv2051c)^28^ then acylates the free amine of the now N-terminal Cys, yielding the mature, triacylated lipoprotein. All three enzymes are essential for *Mtb* survival^17–20^.

**Figure 1.**
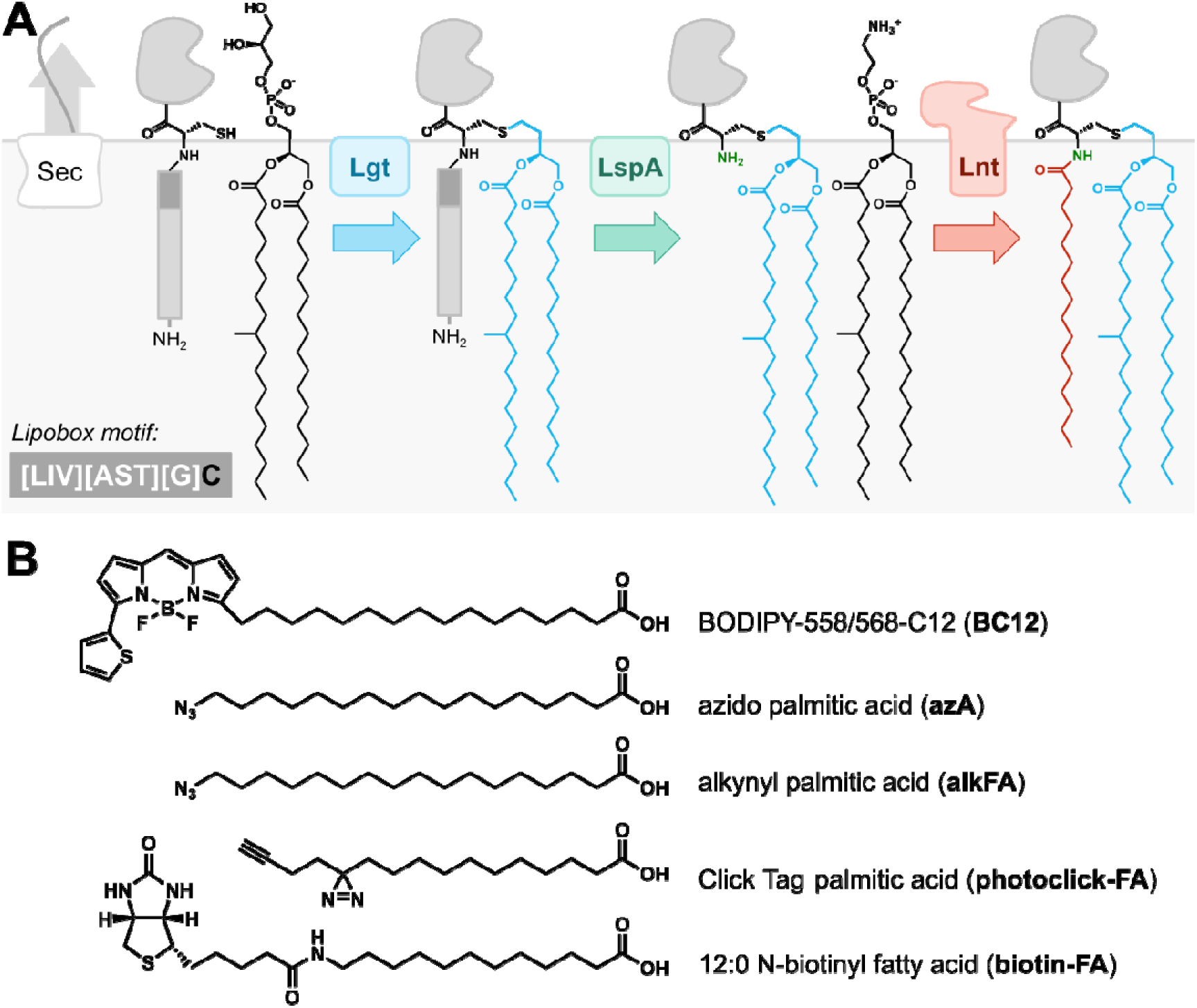
Fatty acids are incorporated into cell envelope proteins via the lipoprotein processing pathway. A) Following export, primarily by the Sec secretion system, proteins are targeted for lipoprotein processing via a lipobox motif (dark grey fill) containing a conserved cysteine. In the first committed step, Lgt modifies the Cys sidechain with a diacylglyceryl group via a thioether linkage (blue). LspA then cleaves the signal peptide, leaving the Cys as the N-terminal residue (green). Lnt then esterifies the amino terminus with a fatty acid (red), yielding the final N-terminally triacylated protein product. The lipobox motif and phosphatidyl glycerol (for Lgt) and phosphatidyl ethanolamine (for Lnt) donors are presumed based on the canonical bacterial lipoprotein biosynthesis pathway^29^. Fatty acid chains (16:0; 19:0) are shown as reported for PG^30^ and presumed for PE. One of two possible configurations for sn-1/sn-2 is displayed and tuberculostearic acid is depicted as the most likely assignment for 19:0. (Signal peptide, chemical structures, and proteins are not to scale relative to the plasma membrane.) B) Structures of modified fatty acids used in this study with commercial names and abbreviations (bold).

Interestingly, although the final enzyme Lnt is present in mycobacteria, diacylated forms are still detected, raising the question of whether these intermediates are functionally distinct from the mature, triacylated forms^28,30^. Also, *Mtb* Lnt activity lies within a bifunctional enzyme with two domains; the other domain is a polyprenol-monophosphomannose synthase involved in the synthesis of the mycobacterial lipoglycans lipomannan and lipoarabinomannan^31^. The possible connection between lipoprotein modification and lipid synthesis remains unexplored.

Lipoproteins have diverse functions in the cell envelope, such as metabolite transport and cell envelope synthesis; they have also been associated with virulence and host defense^23^. Of the 116 proteins most recently catalogued as lipoproteins, 78 have functional annotations, although many of these are only predictive^23^. Regarding the need for lipid modification, one hypothesis is that it serves as localization signal that facilitates function, for example, by positioning proteins proximal to membranes. Despite the diverse and critical roles lipoproteins play in *Mtb*, they remain poorly characterized with respect to their lipid modifications, with few experimentally verified exceptions. Overall, many mysteries remain regarding lipoprotein function and metabolism.

Finally, while other diderm bacteria such as *E. coli* transport the majority of their lipoproteins to the outer membrane using the Lol pathway^32^, mycobacteria lack direct Lol homologues, consistent with their closer phylogenetic relationship to monoderm gram positives^33^. Whether mycobacteria transport lipoproteins beyond the plasma membrane remains an open question.

Experimental validation and characterization of lipoproteins has relied on metabolic labeling, mass spectrometric analysis, or a combination of bioinformatics and subcellular fractionation. A lipobox motif and detection in the membrane fraction^34–37^ provide indirect evidence for individual lipoproteins. We recently experimentally confirmed the cell wall localization of 49 predicted lipoproteins by compartment-specific proximity labeling using the engineered peroxidase APEX2^13^. Post-translational lipid modification of the mycobacterial lipoproteins LpqH, LpqL, LprF, and LppX has also been characterized by mass spectrometry^30,38^. Incorporation of ^3^H- or ^14^C-fatty acids affords more direct evidence for the addition of the lipid modification, as has been applied to LpqH^39^ and LppX^40^, but is less broadly accessible due to expense and safety and handling requirements for radioisotopes. Finally, none of these methods enable further downstream applications such as enrichment and identification.

We posited that fatty acid analogues that are tolerated by the lipoprotein synthesis machinery and enable other modes of detection would facilitate detailed study of individual lipoproteins and the lipoprotein proteome as a whole. Several modified fatty acids have been used to track fatty acid uptake and metabolism in mycobacteria, including in the context of infection, most notably those modified by the fluorophore BODIPY. THP-1-derived macrophage-like cells metabolize BODIPY C12 into triacylglycerides (TAGs), which are then taken up by intracellular *Mtb*^41^. The related BODIPY FL C16 behaves similarly^42^ and has been developed into an assay to quantify incorporation of host-derived BODIPY FL C16 into intracellular *Mtb* by flow cytometry^43^. Mycobacteria in ex vivo infection and in vitro culture incorporate BODIPY fatty acids and/or their metabolites into intracellular lipid inclusions ^20,21,23^. Also, uptake of BODIPY FL C16 is dependent on proteins that stabilize the Mce1 and Mce4 transporter systems^42^. While these data support incorporation of fatty acid analogues into lipids, one study showed that *M. smegmatis* (*Msm*), a fast-growing non-pathogenic relative of *Mtb*, incorporated an alkynyl fatty acid into proteins^44^. The related species *Corynebacterium glutamicum*, which shares overall cell envelope architecture with mycobacteria, incorporates acyltrehalose and fatty acid analogues, including a similar medium-chain alkynyl fatty acid, into mycolic acid modifications on proteins, providing labeling tools to characterize this post-translational modification^45–48^; there have not yet been reports of mycoloylated proteins in mycobacteria such as *Msm*. Overall, fatty acid analogues show promise as labels for tracking lipid-modified metabolites, but their fates in mycobacteria have not been characterized.

In this work we explored the potential of BODIPY-modified and other fatty acid analogues to label proteins in mycobacteria. We found that a BODIPY fatty acid readily labels proteins in both *Msm* and *Mtb* and thus provides a facile alternative to radiolabeling. We further showed that fatty acids containing functional groups compatible with click chemistry modification were also incorporated into proteins and enabled additional modes of detection, with the potential for downstream enrichment and identification. Labeling by an azido-fatty acid was dependent on an enzyme in the lipoprotein biosynthesis pathway and was labile to base hydrolysis, supporting that modified proteins are lipoproteins. Using methods that selectively reveal surface-associated proteins, we found evidence that lipoproteins are exposed at the cell surface, suggesting the transport of lipoproteins beyond the plasma membrane in mycobacteria. Overall, we validated a toolbox of readily available probes for tracking lipid-modified proteins that are expected to accelerate research into mycobacterial lipoprotein identification, synthesis, localization, and, most notably, transport.

## RESULTS AND DISCUSSION

### Mycobacteria metabolically incorporate a variety of fatty acid analogues into proteins

We first confirmed that the fluorescent fatty acid analogue BODIPY 558/568 C12 (BC12; **Figure 1B**) is directly metabolized by mycobacteria. Based on previous studies, we hypothesized that BC12 would be incorporated by mycobacteria into lipids such as TAGs. When cultured in the presence of BC12, *M. smegmatis* (*Msm*), a fast-growing, non-pathogenetic relative of *M. tuberculosis*, rapidly generated apolar lipids as the dominant fluorescent species, based on thin-layer chromatography analysis of solvent extracts and comparison to a standard generated by feeding BC12 to THP-1-derived macrophage-like cells (**Figure S1**).

Rangan et al. previously showed that another fatty acid analogue, alkynyl-fatty acid (alkFA; specifically, chain-length variant alk-14) labels proteins in multiple bacterial species, including *Msm*^44^. We correspondingly confirmed and further characterized protein modification by BC12. Fluorescence associated with proteins was BC12 dose-dependent and incorporation was abrogated in heat-killed cells, showing that incorporation is an active process and that the observed signal is not due to non-covalent association of the label with proteins (**Figure 2A**). *Mtb* also incorporated BC12 into proteins and this signal was suppressed in both *Msm* and *Mtb* by the addition of OADC, a common growth medium supplement that includes albumin (0.5% w/v final concentration) and oleic acid (177 µM compared to 5 µM BC12) (**Figure 2A, B**). Addition of ADC, which contains albumin but not oleic acid, similarly suppressed signal in *Msm*, indicating that sequestration by binding to albumin is a major factor in blocking BC12 incorporation into proteins (**Figure S2**).

**Figure 2.**
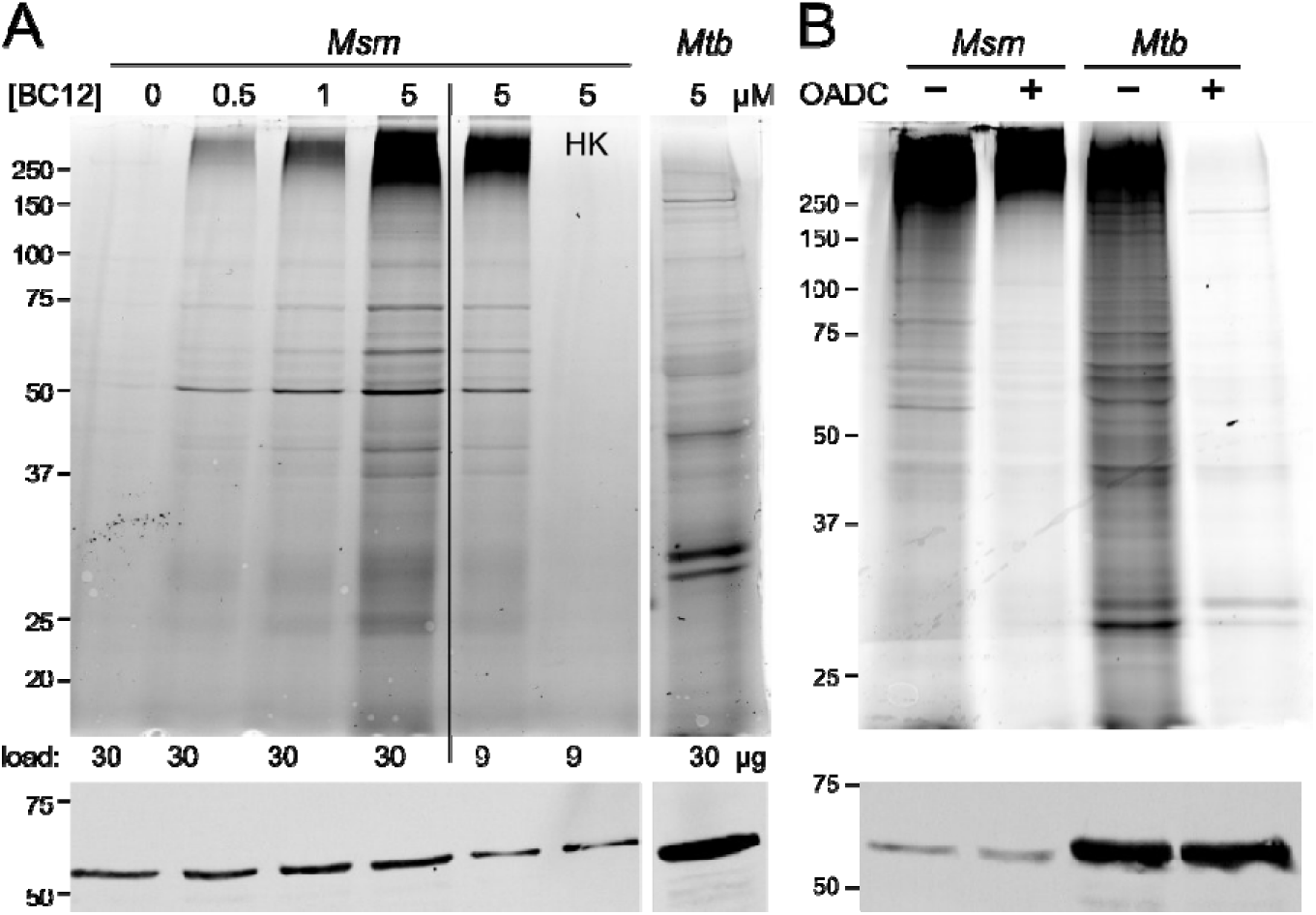
*Msm* and *Mtb* incorporate the modified fatty acid BC12 into proteins. A) *Msm* or *Mtb* was incubated with varying concentrations of BODIPY-C12 (BC12) for ∼1 doubling time (2 h for *Msm*, 20 h for *Mtb*). Five µM BC12 was used for all subsequent experiments. Heat-killed (HK) cells served as a negative control and total lysates were analyzed by SDS-PAGE. B) Same as in A) except −/+ 10% OADC was supplemented during BC12 incubation.All samples in each panel were run on the same gel; lanes scanned in separate detection channels appear slightly separated. GroEL immunoblot was used as a loading control. Data are representative of at least n = 3 independent experiments.

Overall, these data supported BC12 feeding as a facile, accessible method to detect lipid modifications on proteins. We then tested other modified fatty acid that would enable expanded opportunities for detection and affinity enrichment. We tested fatty acids modified with biotin or with an azide group that allows biorthogonal “click” coupling to a wide range of commercially available reagents (**Figure 1B, S3**). Incubating *Msm* with 12:0 N-biotinyl fatty acid (biotin-FA) did not yield any biotin-FA-dependent protein biotinylation (**Figure S4**). In contrast, incubation with alkFA followed by coupling to az-AZ488 also yielded protein labeling, as previously reported^44^ (**Figure 5**). Similarly, azido palmitic acid (azFA) was readily incorporated into *Msm* proteins, as detected after subjecting total lysates to copper-dependent azide-alkyne coupling (CuAAC) to a fluorescent alkyne (alk- Z488) (**Figure 3A**). In contrast, treatment of lysates with dibenzocyclooctyne (DBCO) reagents (DBCO-PEG4-biotin, DBCO-AF488) for strain-promoted azide-alkyne coupling (SPAAC) led to abundant signal that was not dependent on azFA (**Figure S4B,C**). This is likely due to azide-independent coupling to free thiols on cysteines^49^. Overall, these results support CuAAC modification of incorporated azFA and alkFA as the most versatile approach for tagging lipid-modified proteins.

**Figure 3.**
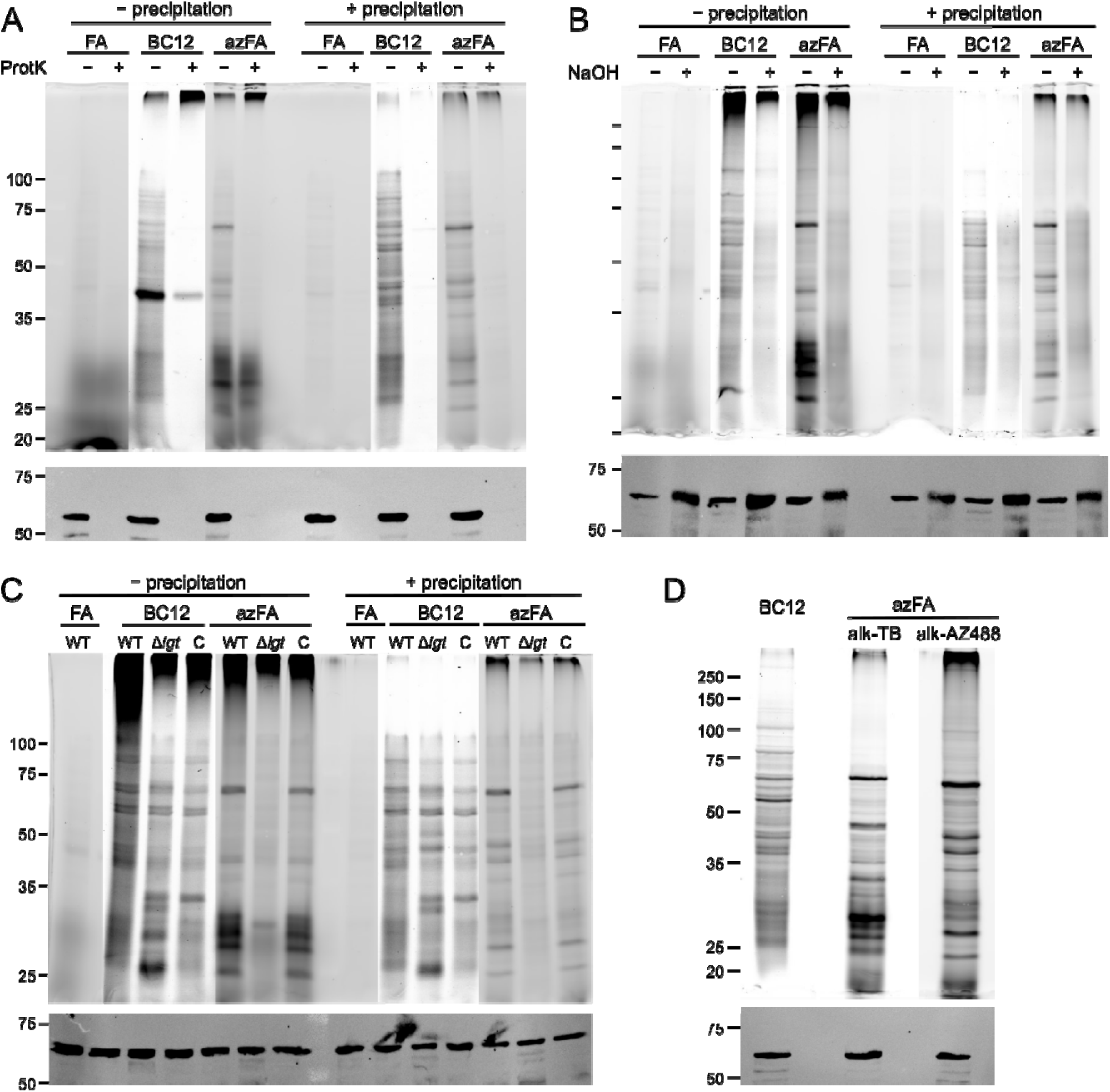
Modified fatty acids added to proteins are labile to base hydrolysis with some dependence on the Lgt MSMEG_3222 for incorporation. *Msm* (A-D) wild type (“WT”), (C) Δ*MSMEG_3222* null (“Δ*lgt*”) or Δ*MSMEG_3222*::*MSMEG_3222* complement (“C”) strains were incubated with 5 µM BC12, 20 µM azFA, or 20 µM palmitic acid as a negative control (“FA”) for 2 h before harvesting. Total lysates of azFA-treated cells were subjected to CuAAC with alk-AZ488 or alk-TB. Lysates were then treated with (A) 0.05 mg/mL proteinase K or (B) 0.1 M sodium hydroxide for 1h before inactivation or quenching and protein precipitation. All samples in each panel were run on the same gel; some lanes were scanned in separate detection channels. GroEL immunoblot was used as a loading control. Data are representative of n = 1-3 independent experiments.

Given that fatty acids are incorporated into lipids, we noted that species like lipoglycans lipomannan and lipoarabinomannan also migrate by SDS-PAGE and thus could contribute to the observed signal. Therefore, we tested that the observed signal is due to labeled proteins by removing lipids by protein precipitation and by treating lysates with protease and. Precipitation preserved the majority of the observed signal, but with the selective loss of diffuse bands in the 25-35 kDa region, where lipomannan and lipoarabinomannan species migrate (**Figure 3**). The addition of proteinase K reduced signal from both BC12- and azFA-treated samples (**Figure 3A**). In a separate experiment, treatment with sodium hydroxide showed that the modifications were base-labile (**Figure 3B**). These results supported protein esterification and thus suggested modification via the lipoprotein biosynthesis pathway.

We sought to test first in *Mtb* whether protein modification was indeed dependent on lipoprotein biosynthesis. Since the lipoprotein processing enzymes Lgt, LspA, and Lnt are all essential in *Mtb*, we used inducible CRISPRi^50^ to knock down expression of the first enzyme in the pathway, Lgt*. Lgt* was essential for *Mtb* survival on agar when a guide RNA targeting *lgt* (*rv1614*) was induced with anhydrotetracycline (ATc) (**Figure S5A**). However, in liquid culture, prolonged ATc treatment only moderately compromised survival or reduced RNA levels (**Figure S5B-D**). Also, ATc treatment did not affect metabolic labeling of proteins by BC12, even under conditions that produced the most pronounced growth defect (**Figure S5E**). These results are consistent with a CRISPRi screen in *Mtb* showing that *lgt* is a relatively insensitive target, but did not provide conclusive evidence regarding the involvement of Lgt in incorporating BC12 into proteins.

We then turned to *Msm*, which encodes two *lgt* homologues, *MSMEG_3222* and *MSMEG_5408*. Neither is required for *Msm* survival, suggesting that they have redundant function. *MSMEG_3222* has higher homology to *Mtb lgt* and a previous study showed that a targeted deletion strain Δ*MSMEG_3222* exhibited reduced incorporation of ^14^C-palmitic acid into proteins and altered localization of known lipoproteins compared to the wild type, supporting a role for MSMEG_3222 in lipoprotein biosynthesis and its assignment as an Lgt^26^. To test whether *MSMEG_3222* is required for modified fatty acid incorporation, we constructed *Msm* Δ*MSMEG_3222* by recombineering (**Figure S6**) and found that loss of MSMEG_3222 led to a reduction in protein-associated azFA signal and this could be complemented (**Figure 3C**). In contrast, the incorporation of BC12 was not obviously affected (**Figure 3C**). This suggests that BC12 is added to proteins via a pathway that does not involve MSMEG_3222, as addressed further in the Discussion. We also noted that BC12 and azFA labeling yielded distinct protein patterns (**Figure 3D**). We hypothesized that these differences do not necessarily indicate that BC12 and azFA are incorporated into different proteins, but rather, arise from the chemical properties of the modified fatty acid, including the nature of the click reagent conjugated to the azide (**Figure 1B, S3**). The distinct migration of azFA-labeled proteins when coupled to alk-AZ488 vs. tetramethylrhodamine (TAMRA)-biotin-alkyne (alk-TB) supported this idea (**Figure 3D**). Overall, while the precise nature of BC12 protein esterification remains to be determined, the data are consistent with the addition of azFA as a lipoprotein modification.

Our further characterization efforts thus focused on azFA. To further confirm its incorporation into lipoproteins, we tested integration into the known lipoprotein LprG. *Msm* expressing the *Msm* LprG homologue (MSMEG_3070) with a C-terminal 3xFLAG tag was incubated with azFA and total lysates were subjected to CuAAC to alk-TB. Alk-TB includes tetramethylrhodamine (TAMRA) for fluorescence visualization and biotin for affinity enrichment (**Figure S3**). Similar to alk-AZ488 labeling, detection of fluorescence (**Figure 4A**) and biotinylation (**Figure S7**) confirmed the specificity of azFA labeling (+ azFA) vs. palmitic acid (− azFA) controls. Following avidin affinity enrichment, *Msm* LprG- 3xFLAG was detected only from cells treated with azFA (**Figure 4B**). The migration of the enriched subpopulation was slower than that of the dominant form in the input, which is presumed to be the mature triacylated form. If this is the case, then these data further support our earlier hypothesis that modified fatty acids alter the migration of lipoproteins (**Figure 3D**). This is similar to our observation of altered LprG migration after modification with biotin by proximity labeling^51^. In summary, these results show that BC12 and azFA are esterified onto proteins and that azFA incorporation depends on lipoprotein biosynthesis.

**Figure 4.**
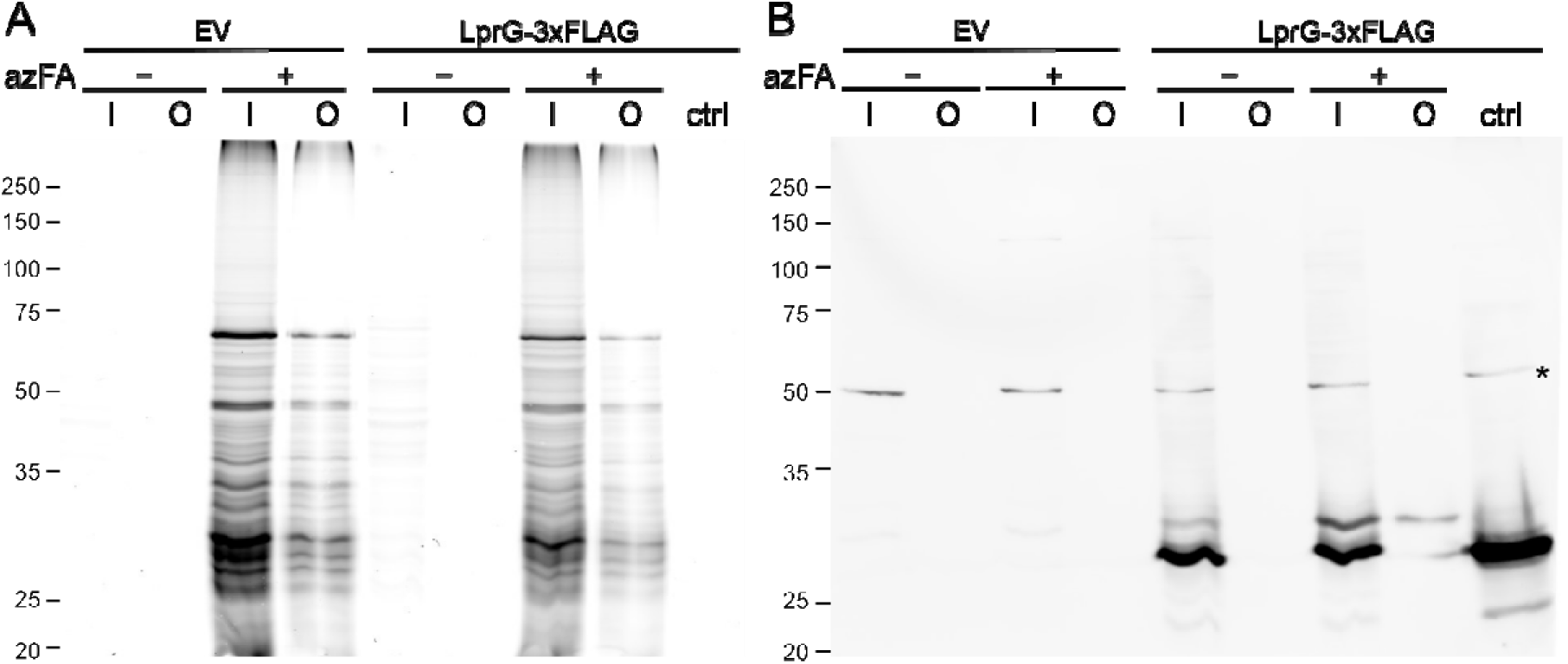
Enrichment of azFA-labeled proteins confirms modification of the lipoprotein LprG. *Msm* expressing LprG-3xFLAG (MSMEG_3070-3xFLAG) or harboring empty vector control (EV) was cultured with azFA for 2 h. Total lysates were subjected to CuAAC with alk-TB followed by avidin enrichment. (A) Fluorescence scan for tetramethylrhodamine (TAMRA) and (B) anti-FLAG immunoblot. (I = input, O = output, ctrl = untreated total lysate from *Msm* LprG-3xFLAG as anti-FLAG positive control. Asterisk indicates non-specific cross-reacting protein.)

### Lipoproteins are associated with the mycobacterial cell surface

As noted in the Introduction, most gram-negative lipoproteins are targeted to the outer membrane and many are found at the cell surface^52^. However, the localization of lipoproteins within the mycobacterial cell envelope has remained an open question. To address this, we used several independent approaches based on azFA labeling to test whether mycobacterial lipoproteins are surface accessible.

First, mycobacteria are commonly cultured in the presence of Tween 80 detergent to disperse cells, but in the absence of detergent, they retain more surface-associated components such as carbohydrates and proteins, which can be subsequently extracted with Tween 80 for analysis^53^. *Msm* was cultured in medium with and without detergent and then incubated with azFA to label lipoproteins. Cells were then harvested and washed with detergent buffer to generate a Tween 80 extract. The remaining cell pellet was then lysed to obtain a total lysate. Both Tween extracts and total lysates were subjected to CuAAC to alk-AZ488 While the total lysates yielded similar results, only Tween extracts from cellscultured without detergent yielded labeled proteins and this profile was distinct from that of total lyates (**Figure 5A**). This result indicates that a subset of lipoproteins was removed by gentle detergent treatment, with the implication that these proteins were at the cell surface.

**Figure 5.**
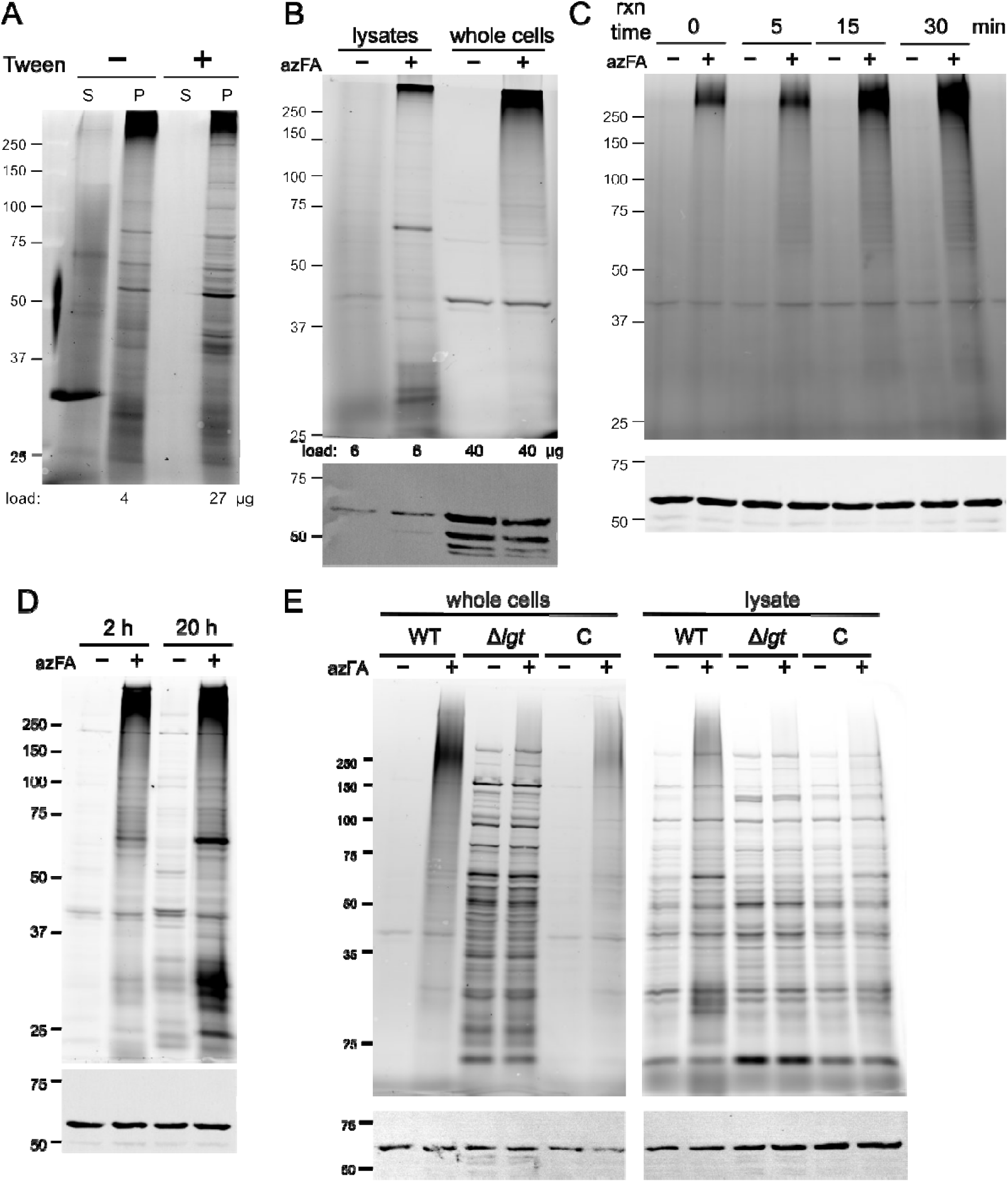
Click coupling to detergent extracts and whole-cells reveals surface-accessible lipoproteins. A) *Msm* was cultured without (−) or with (+) Tween 80, incubated with 20 µM azFA for 2 h. Cells were then extracted with 1% Tween 80. The resulting supernatant extract (S) and the total lysate of the remaining cell pellet (P) were subjected to CuAAC. Protein load for −/+ Tween pellet fractions were normalized approximately for the degree of observed labeling. B) *Msm* in standard +Tween medium was incubated with µM for 2 h. CuAAC to alk-AZ488 was then performed on whole cells and total lysates. Protein load for lysate and whole-cell samples were normalized approximately for the degree of observed labeling. (C, D) Same as in B) except azFA was incubated with *Msm* for the specified times prior to CuAAC to alk-AZ488 on whole cells. E) Same as in B) using *Msm* wild- type, Δ*MSMEG_3222* (Δ*lgt*), and complement Δ*MSMEG_3222::MSMEG_3222* (C) strains.

**Figure 6.**
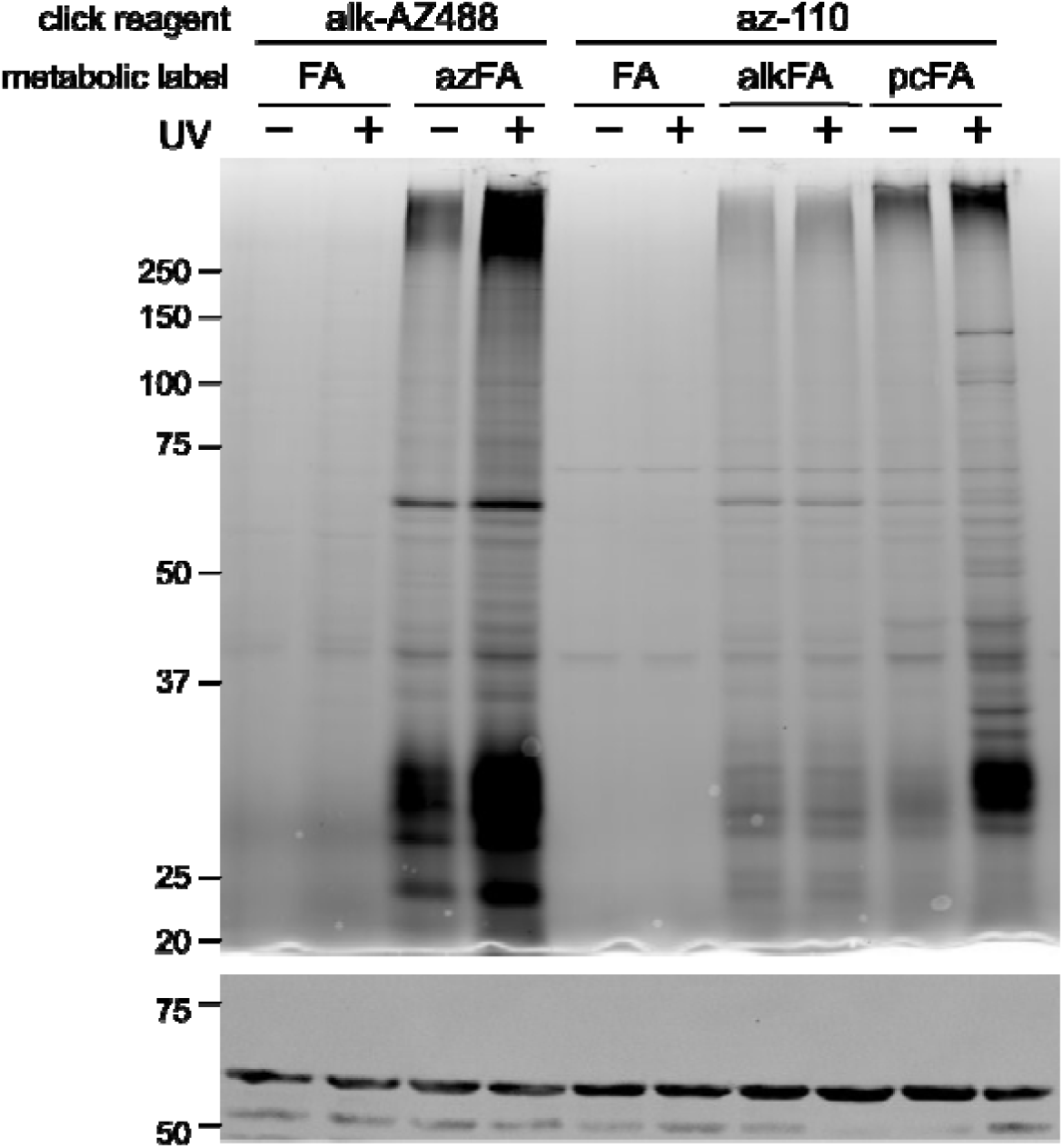
Incorporation of a photoclick-FA in *Msm* yields UV-dependent adducts. *Msm* was incubated with palmitic acid (FA), azFA, alkFA, or pcFA for 30 min. and then UV irradiated for a total of 30 min. Total lysates were subjected to CuAAC with either alk-AZ488 or az-110 as indicated. GroEL immunoblot served as a loading control. Data are representative of n = 3 independent experiments.

Next, we hypothesized that the negative charges of the two sulfonic acid groups on AZDye 488 (**Figure S3**) would retard alk-AZ488 permeation into whole cells, since measurements of *Msm* zeta potential support a net negative charge^54,55^.We predicted that performing CuAAC with alk-AZ488 on whole cells labeled with azFA would selectively label surface-accessible lipoproteins. This indeed yielded azFA-dependent protein labeling with a pattern distinct from that of similarly treated total lysates (**Figure 5B**).

Finally, whole cells subjected to SPAAC by incubation with DBCO-AZ488 showed azFA- and time-dependent labeling (0-30 min; **Figure 5C**), in contrast to total lysates treated the same way (**Figure S2B, C**). Importantly, longer incubation times (2 and 20 h) yielded labeling reminiscent of lysates treated with DBCO (**Figure 5D**, **Figure S3B-C**) and this labeling was not fully azFA- dependent. These results support surface-specific tagging by SPAAC at shorter incubation times, with eventual intracellular accumulation of DBCO-AZ488.

We further reasoned that the lipoprotein-deficient strain Δ*MSMEG_3222* would have reduced levels of surface-accessible lipoproteins and tested this hypothesis by incubating wild-type, Δ*MSMEG_3222* and complement strains with azFA followed by treatment of whole cells with DBCO- AZ488 (**Figure 5E**). Wild-type and complement strains indeed displayed similar patterns of azFA- dependent labeling. However, rather than showing reduced labeling, Δ*MSMEG_3222* yielded azFA- independent labeling similar to that of lysates by DBCO reagents (**Figure 5E**, **Figure S2B-C**). This suggests that DBCO-AZ488 accumulates intracellularly more quickly in Δ*MSMEG_3222* than the wild type and thus, that loss of *MSMEG_3222* compromises the cell envelope. However, this experiment was not conclusive for changes in surface-accessible lipoproteins upon a reduction in Lgt activity.

Overall, these multiple lines of evidence support the conclusion that mycobacteria display lipoproteins at the cell surface, including in a loosely associated layer that is not present under standard culture conditions with detergent in the growth medium.

### A photocrosslinking fatty acid enables the detection of lipid-protein interactions

We posited that metabolic labeling would also afford an opportunity to detect lipid-protein interactions using a fatty acid modified with a photocrosslinking group. *Msm* was incubated with a palmitic acid analogue containing a terminal alkyne and a photoactivatable diazirine crosslinker (photoclick-FA)^56^ and UV-irradiated prior to lysis and CuAAC to a fluorescent azide (az-AZ488). In the absence of UV irradiation, feeding with photoclick-FA revealed a similar protein labeling pattern (**Figure 5**) to that of other click-modified fatty acid coupled to AZ488s, including azFA and alkFA (**Figure 2, 4**). However, only treatment with photoclick-FA yielded additional, higher-molecular weightprotein bands after UV irradiation, consistent with the formation of lipoprotein-protein adducts due to crosslinking of the lipid modification to an interacting protein (**Figure 5**).

## DISCUSSION

Here we have expanded and validated metabolic labeling with modified fatty acids as a method to track lipid-modified proteins in mycobacteria. We found that *Msm* and *Mtb* incorporated into proteins a range of fatty acid derivatives, including those modified with fluorophores, click chemistry-compatible groups, and/or photo-crosslinking groups. Of all the modified fatty acids tested, only biotin-FA was not clearly incorporated into proteins. Lack of detection could be due to limited intracellular accumulation biotin-FA on the 2-hour timescale of incubation, although we have found that other alkyl biotin derivatives, such as protein-reactive biotin-NHS esters and the proximity labeling substrate biotin-phenol, show evidence of accumulation (via protein modification) within 30 min^13,51^. Another possibility is that the biotin moiety is significantly metabolized during the incubation, preventing detectable incorporation of the modified fatty acid into proteins.

Base lability, dependence on Lgt, and incorporation in the known lipoprotein LprG indicate that proteins are modified via the lipoprotein pathway (**Figure 1**), at least in the case of azFA. In contrast, BC12 labeling does not strongly depend on the Lgt homologue MSMEG_3222 in *Msm*. The loss of MSMEG_3222 does not completely abrogate azFA labeling, suggesting redundancy with the other *lgt* homologue *MSMEG_5408*. PG is considered to be the absolute substrate for other bacterial Lgts based on its lack of activity with PE or cardiolipin *in vitro*^57^, but it is possible that mycobacterial Lgt enzymes have a broader substrate range. Furthermore, MSMEG_5408 may have a different lipid substrate preference than MSMEG_3222. We speculate that BC12 is preferentially added to lipids that are processed more readily by MSMEG_5408 and that thus its addition to proteins less dependent on MSMEG_3222. Alternatively, mycobacteria may add BC12 not only to glycerolipids, but also ultra-long fatty acids known as mycolic acids, via fatty acid synthase II (FAS-II). In *Corynebacterium glutamicum*, mycolic acids are esterified onto proteins by the acyltransferase Cmt1 (also known as CMytC). However, neither a corresponding enzyme nor evidence of protein O-mycoloylation has yet been reported in mycobacteria. A full elucidation of protein modification with BC12 thus awaits further studies.

The incorporation of fatty acids that allow downstream enrichment of labeled proteins will enable not only the confirmation of specific lipoproteins, as demonstrated here for LprG, but also the identification of the lipoproteome, as a goal for future studies. Most importantly, our results showed that lipoproteins are surface accessible and thereby indicate that mycobacteria have a transport pathway for moving lipoproteins from the plasma membrane to the mycomembrane. While the gram-negative Lol transport pathway has no homologues in mycobacteria, the soluble protein LolA and the lipoprotein LolB have strong structural homology to the lipid-binding lipoprotein (Llp) family in mycobacteria (LprG, LppX, LprA, LprF)^33^. All four proteins have been implicated in lipid binding and/or transport, but LprG alone is conserved across mycobacteria. While LprG has so far been implicated in lipid transport, it binds to triacyl lipids^58,59^, consistent with a possible role in lipoprotein transport.

Finally, the presence of both diacyl and triacyl lipoproteins in mycobacteria^26,30^ suggests the tantalizing hypothesis that triacylation serves as a signal for lipoprotein transport across the cell envelope. This would contrast with gram-negative lipoprotein transport in *E. coli*, in which recognition by the transport machinery is encoded by conserved N-terminal residues^60^.

The formation of UV-dependent adducts following labeling with photoclick-FA provides a means to detect interacting proteins. Given that BC12 is also metabolized into lipids (**Figure S1**), photoclick-FA may similarly yield labeled lipids that then crosslink to proteins. This approach offers an additional opportunity identify lipid-interacting proteins, as has been done with analogous trehalose monomycolate lipid analogues. By the same token, using this method to discover proteins that interact specifically with lipoproteins will require careful validation. Overall, our results characterizing the metabolic labeling of protein lipid modifications pave the way for the broader use of this method to track the biosynthesis, inhibition, and binding of lipoproteins not only in mycobacteria, but also in other bacteria in which these processes are relatively poorly understood.

## METHODS

### Bacterial strains and growth

All strains, oligonucleotides and plasmids used in this study are detailed in Tables S1-S3. *Mycobacterium smegmatis* mc^2^155 (*Msm*; ATCC 700084) was cultured in Middlebrook 7H9 medium (HI MEDIA) with 1% w/v casamino acids (VWR), 0.2% w/v glucose, 0.05% v/v Tween 80 at 37 °C with shaking at 250 rpm unless otherwise noted. For growth on solid medium, *Msm* was propagated on Middlebrook 7H11 agar with 10% v/v albumin-dextrose-catalase (ADC) supplement (BD), 0.5% v/v glycerol, 0.05% v/v Tween 80. *M. tuberculosis* H37Rv Δ*lysA* Δ*panCD* (*Mtb*; gift of William Jacobs)^61^ was cultured in Middlebrook 7H9 with 10% v/v oleic acid-albumin-dextrose-catalase (OADC) supplement (BD), 0.5% v/v glycerol, 0.2% w/v casamino acids, 0.025% v/v Tyloxapol and 80 µg/mL lysine and 24 µg/mL pantothenate or on Middlebrook 7H10 agar with the same supplements except without Tyloxapol. All cultures were incubated at 37 °C with shaking at 110 rpm. Bacteria were selected on kanamycin (25 µg/mL) or hygromycin (50 µg/mL) as appropriate (Table S3). Cells were harvested by centrifugation at 4,000 x g for 5 min and all steps were performed at 22 °C unless otherwise indicated.

### Generation and confirmation of *Msm* Δ*MSMEG_3222*

A targeted knockout strain of *MSMEG_3222* was generated in *Msm* as previously reported^62^. Briefly, 5’ (321 bp) and 3’ (248 bp) sequences flanking *MSMEG_3222* were cloned on either side of a hygromycin resistance cassette via the HindIII and XbaI sites of pJSC407 (gift of Jeffrey Cox) using In-Fusion Cloning (Takara Bio). The resulting plasmid was sequence confirmed and used as a template for PCR using oligonucleotides omlp1085 and omlp1088 (Table S2) to generate a recombineering substrate that was transformed directly into *Msm::*pNIT-RecET expressing the recombinase following treatment with isovaleronitrile. Successful recombinants were selected on agar containing hygromycin and individual clones confirmed by PCR (Figure S1) and sequencing. A complement plasmid was generated by cloning *MSMEG_3222* (including 998 nt 5’ of the start codon as a native promoter) into the integrating plasmid pMV306 via the XbaI and ClaI sites using In-Fusion Cloning. The resulting plasmid was sequence confirmed, transformed into Δ*MSMEG_3222* and selected on agar containing kanamycin.

### Metabolic incorporation of fatty acids in *Msm*

*Msm* was cultured to mid-logarithmic phase (OD_600_ ∼1.5) and then diluted to OD_600_ 0.5 and incubated for 2 h (unless otherwise specified) at 37 °C with palmitate as a negative control (FA; Acros Organics), BODIPY 558/568 C12 (BC12; ThermoFisher, cat # D3835), azido palmitic acid (azFA; Vector Laboratories, cat # CCT-1246), alkynyl palmitic acid (alkFA; Vector Laboratories, cat # CCT-1165), Click Tag™ palmitic acid (photoclick-FA; Cayman Chemical, cat # 37969), or 12:0 N-biotinyl fatty acid (biotin-FA; Avanti Lipids, cat # 860557) at the final concentrations indicated in each figure (0-20 µM; **Figure S3**). For *Mtb,* initial cultures were grown to OD_600_ 1 and then diluted to OD_600_ 0.5 in 20 mL of culture medium containing 5 µM BC12. Cultures were harvested after 1 doubling time (∼16 hours). All fatty acids were dissolved in DMSO and final DMSO concentrations were 0.6% for 12:0 N-biotinyl fatty acid and 0.05-0.1% for all others.

Experiments involving total lysates resulting from BC12 or azFA treatment were derived from 10 mL of bacterial culture per condition except where noted. Experiments with photoclick-FA used 30- 40 mL of bacterial culture per condition. After incubation with fatty acid, cells were harvested by centrifugation and washed three times with an equal volume of PBS with 0.05% v/v Tween 80 (PBST). Pellets were processed immediately or stored at -80 °C until further use.

### Detergent extraction of whole cells

*Msm* was cultured to mid-logarithmic phase (OD_600_ ∼1.5) and then subcultured to OD_600_ ∼0.02 in 10 mL culture medium with or without 0.05% Tween 80 and grown overnight to a final OD_600_ ∼1.5, as determined from the dispersed culture containing Tween 80. Both cultures were diluted to OD_600_ 0.5 and incubated with 5 µM BC12 for 2 h at 37 °C. Harvested cells were washed twice with an equal volume of PBS. The final cell pellet was resuspended in 800 µL PBS with 1% v/v Tween 80. The suspension was transferred to a 1.5-mL microfuge tube and incubated for 30 min with gentle mixing at 22 °C on an orbital shaker. Cells were harvested by centrifugation at 10,000 x *g* for 10 min and 500 µL supernatant was collected as the detergent extract. The remaining cell pellet was processed for total lysate.

### UV irradiation for photo-crosslinking

After incubation with photoclick-FA, cells (corresponding to 30-40 mL culture; see section “Metabolic labeling with fatty acids”) were harvested and resuspended in 2 mL PBST. One 1-mL aliquot was pipetted into a 6-well multiwell plate; the other was kept on ice as the no-irradiation (-UV) control. The multiwell plate was placed on ice and irradiated from above with a UV transilluminator for a total of 30 min with gentle manual agitation every 5 min.

### Preparation of total lysates

Each cell pellet was resuspended in 800-100 µL PBS and added to a 2-mL microfuge tube containing ∼0.5 mL 0.1-mm zirconia beads. Cells were lysed at 6 m/s for 30 s (Bead Ruptor 12, Omni International) for a total of 4 cycles with samples placed on ice for 5 min between cycles. Lysates were clarified by centrifugation at 10,000 x *g* at 4 °C for 10 min and the clarified supernatants were transferred to fresh tubes. *Mtb* cells were subjected to a total of 5 cycles of bead beating and lysates were clarified by centrifugation at 12,000 x *g* for 10 min at 4°C and sterilized through 0.22-µm filters before removal from biosafety level 2 containment for further processing.

### Azide-alkyne coupling

The protein concentrations of total lysates were determined by the BCA assay (Pierce) and normalized to 0.65-1 mg/mL depending on the concentration of the most dilute sample in a given experiment. Copper-mediated azide-alkyne cycloaddition (CuAAC) was performed as previously described^5^. Briefly, reagents were added sequentially to 100 µL normalized total lysate to the following final concentrations: 5 mM sodium ascorbate, 25 µM click reagent, 0.2 mM, 0.2 mM copper sulfate and incubated at 22 °C for 1 h for Figure 1C and 5 min for all other experiments. Reactions were quenched with 1 mM EDTA Click reagents used in this study were AZDye 488 Alkyne (alk-AZ488; Vector Laboratories, cat # CCT-1277), AZDye 488 Azide (az-AZ488; Vector Laboratories, cat # CCT-1275), azido-PEG3-carboxyrhodamine 110 (az-110; ThermoFisher, cat # J65107.MB), and TAMRA Biotin Alkyne (alk-TB, Vector Laboratories cat # CCT-1366) (**Figure S3**).. Where indicated, protein was then precipitated by the sequential addition of 4 volumes of methanol, 1 volume of chloroform, and 3 volumes of water and mixed by vortexing. Samples were centrifuged at 18,000 x *g* for 5 min at 4 °C. After removing the upper aqueous layer, the pellet was washed twice with 3 volumes of methanol and resuspended in 100 µL PBS with 1% SDS. For strain-promoted reactions (SPAAC) with total lysates, 100 µL normalized lysate was incubated with 25 µM AZDye 488 DBCO (DBCO-AZ488; Vector Laboratories, cat # CCT-1278) or Biotin-PEG4-DBCO (DBCO-biotin; Vector Laboratories, cat #OCT-A105) incubated for 1 h at 22 °C with orbital shaking (250 rpm). For SPAAC with whole cells, the cell pellet from 12.5 mL of culture (OD_600_ ∼0.8) was resuspended in 800 µL PBS and incubated with 25 µM DBCO-AZ488 at 22 °C for 0-20 h. Cells were harvested and washed 4 times with an equal volume of PBST before storage or processing for total lysates.

### Protease and alkaline treatments

For protease treatment, proteinase K (Sigma) dissolved at 2.5 mg/mL in 50 mM Tris pH 7.4, 1 mM CaCl_2_, 0.5% SDS was added to quenched CuAAC reactions at a final concentration of 0.05 mg/mL and incubated with orbital shaking (250 rpm) at 37 °C for 1 h.

Proteinase K was deactivated by the addition of 5 mM phenylmethylsulfonylfluoride for 10 min at 22 °C. For alkaline treatment, sodium hydroxide was added directly to quenched CuAAC reactions at a final concentration of 0.1 M and incubated at 37 °C with orbital shaking (250 rpm) for 1 h and then neutralized with a stoichiometric amount of 0.1 M acetic acid. Excess reagent and cleaved label were removed by protein precipitation as above in “Azide-alkyne coupling.”

### Avidin affinity enrichment

*Msm* transformed with empty vector (pGW1-6C)^63^ or an episomal plasmid encoding *Msm* LprG-3xFLAG [pSMT-*Msm*LprG(Y3xH)-3xFLAG-MjtRNA]^64^ were cultured and subjected to azFA treatment as above (“Metabolic incorporation of fatty acids in *Msm*”) in 30-mL cultures. CuAAC with alk-TB was performed as above (“Azide-alkyne coupling”), except reactions were scaled to 400 µg; precipitation steps were performed with centrifugation at 4,000 x *g* for 20 min at 4 °C; and the final protein pellet was resuspended in 500 µL resuspension buffer (0.5% SDS, 0.05% LDAO in water). After reserving 100 µL as input, the remainder was mixed with 116 µL NeutrAvidin agarose beads (pre-equilibrated in resuspension buffer; Pierce/ThermoFisher, cat # 29201) and incubated with rotation at 22 °C for 2 h. Beads were pelleted at 3,000 x *g* for 1 min at 4 °C and washed once each with 2 volumes of resuspension buffer; 8M urea and 1% SDS; and PBS. Bound proteins were eluted by the addition of 126 µL 4X Laemmli sample buffer and boiling at 95 °C for 15 min. For SDS-PAGE, 30 µg input and 6 µL output was analyzed for each sample group.

### Protein analysis by SDS-PAGE

Protein concentrations of clarified lysates (see “Preparation of total lysates” above) were determined by BCA assay (Pierce) and normalized to 0.75 mg/mL with PBS and Laemmli sample loading buffer was added to 1X final concentration. Where applicable, gels were imaged for fluorescence using a Sapphire Bioimager (Azure Biosystems). For immunoblotting, proteins were transferred to nitrocellulose membranes (Trans-Blot Semi-Dry Transfer Cell; Bio-Rad) and probed with α-GroEL (1:5000; Santa Cruz Biotechnology, cat # sc-58170) or α-FLAG M2 (1:1000; Sigma-Aldrich, cat # F1804) and goat anti-mouse IR Dye 800CW (1:15,000, LI-COR, cat # 926−32210). Blots were imaged using an Odyssey CLx imager (LI-COR)Images were analyzed with ImageJ.

## ACKNOWLEDGMENTS

We thank Neetika Jaisinghani for help with experimental design and troubleshooting and for generating the THP-1-derived lipid extract and the J. Seeliger lab for helpful discussions.

