## Supplemental Figures S1-S7 and Supplemental Methods for "Metabolic tagging reveals surface-associated lipoproteins in mycobacteria"

### SUPPLEMENTAL INFORMATION


- **Figure S1.** BODIPY-C12 is metabolically incorporated into *Msm* lipids
- **Figure S2.** Incubation with either ADC or OADC supplement reduces metabolic incorporation of BC12 into proteins in *Msm*
- **Figure S3.** Structures of click reagents used in this study.
- **Figure S4.** *Msm* does not incorporate biotin-palmitic acid into proteins and DBCO reagents label proteins in an azide-independent manner in total lysates.
- **Figure S5.** *M. tuberculosis lgt* is essential but not highly vulnerable by inducible CRISPRi knockdown and suppression does not affect protein labeling by BODIPY-C12
- **Figure S6.** Confirmation of Δ*MSMEG_3222* by PCR and sequencing
- **Figure S7.** *Msm* treatment with azFA followed by CuAAC to alk-TB and detection of biotin confirms azFA incorporation into proteins.
- **Supplemental Methods**

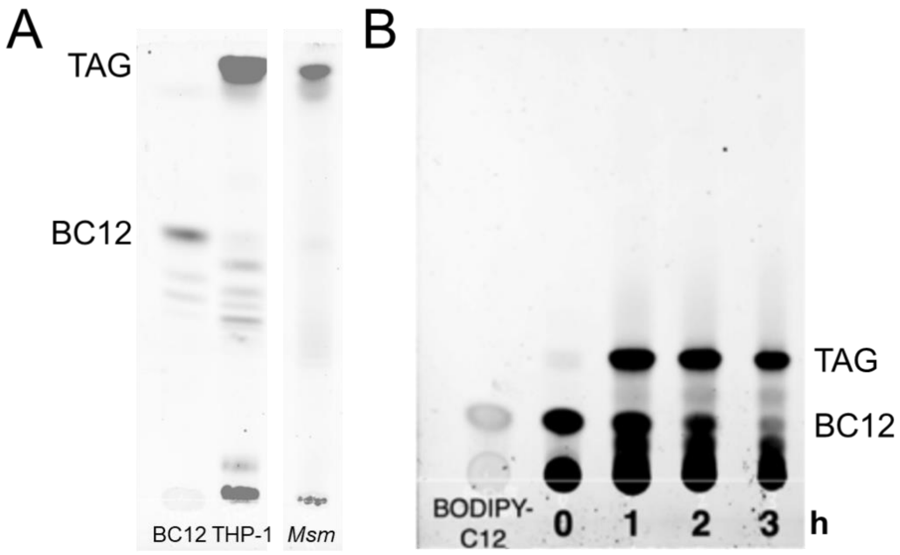


**Figure S1. BODIPY-C12 is metabolically incorporated into *Msm* lipids**

Chloroform:methanol extracts from *Msm* treated with BODIPY 558/568 C12 (BC12) were resolved on silica plates in A) 100:14:0.8 chloroform:methanol:water or B) 70:30:1 hexanes:diethyl ether:acetone mobile phase. In B) extracts were made after the specified times of *Msm* incubation with BC12. Extracts from BC12-treated THP-1 cells served as a migration control for triacylglyerides (TAG) containing BC12 (see Supplemental Methods).

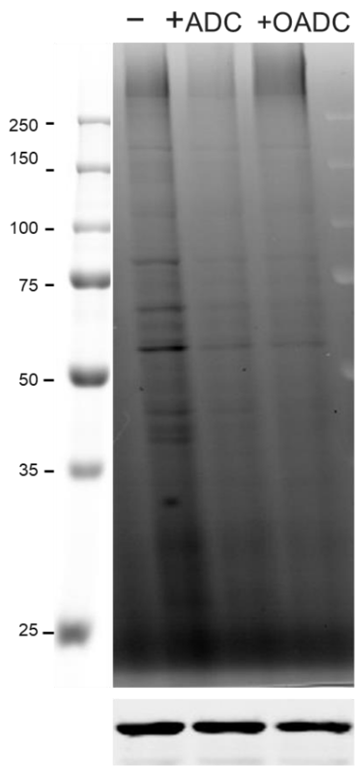


**Figure S2. Incubation with either ADC or OADC supplement reduces metabolic incorporation of BC12 into proteins in *Msm.*** *Msm* was incubated with 5 µM BC12 for 2 h in the culture medium without (−) or with (+) 10% ADC or OADC supplement. GroEL immunoblot served as a loading control. Data are from a single experiment.

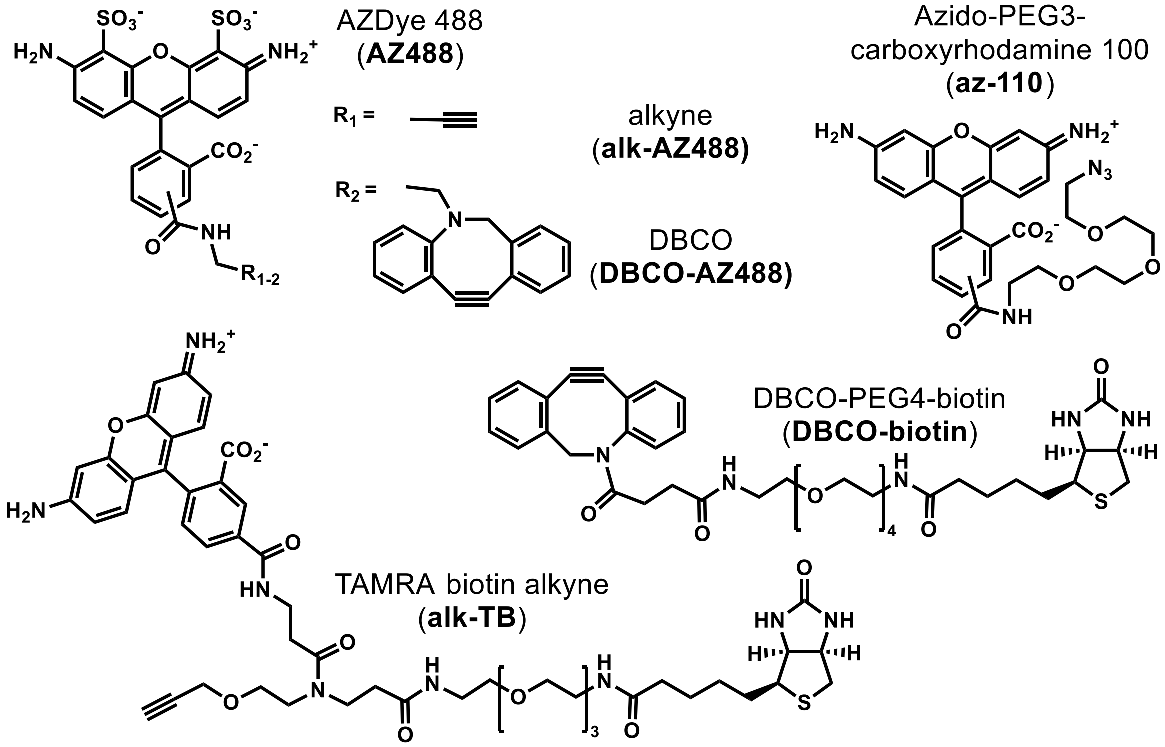


**Figure S3. Structures of click reagents used in this study.** All reagents are commercially available; see main text Methods for details.

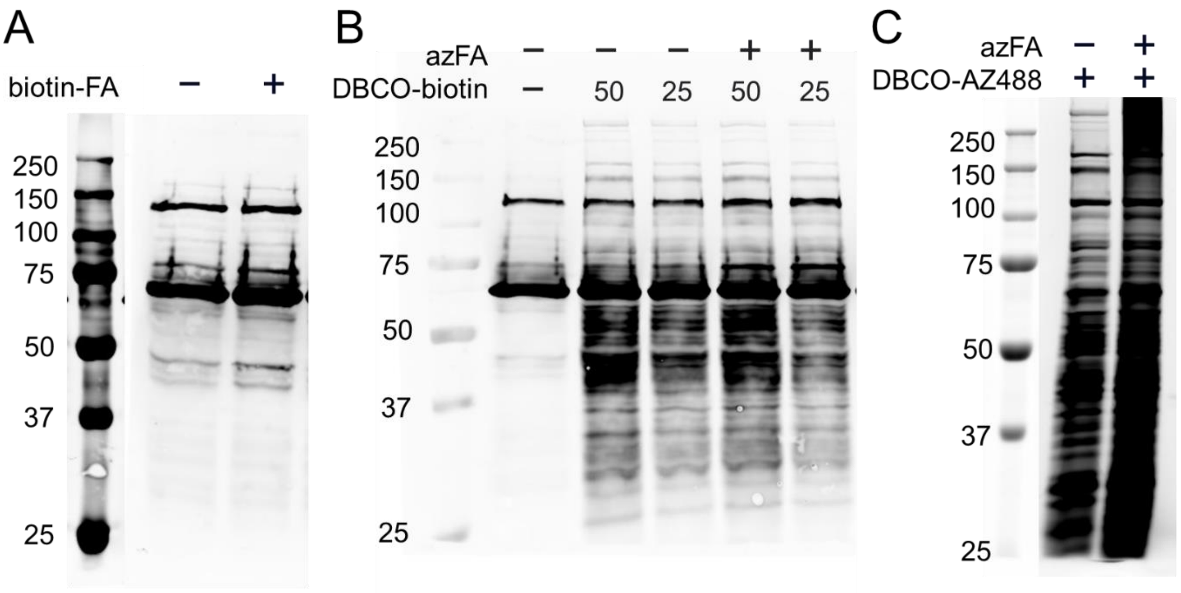


**Figure S4. *Msm* does not incorporate biotin-palmitic acid into proteins and DBCO reagents label proteins in an azide-independent manner.** *Msm* was incubated with 20 µM biotin-palmitic acid (biotin-FA) or azide-palmitic acid (azFA) for 2 h. In B) total lysates were then incubated with the specified DBCO reagents at 25 µM for 30 min. Samples were analyzed by SDS-PAGE and A, B) immunoblot [as described in Methods, with streptavidin IR-Dye 680LT (1:10,000; 926-68031, LI-COR)] or C) fluorescence scanning.

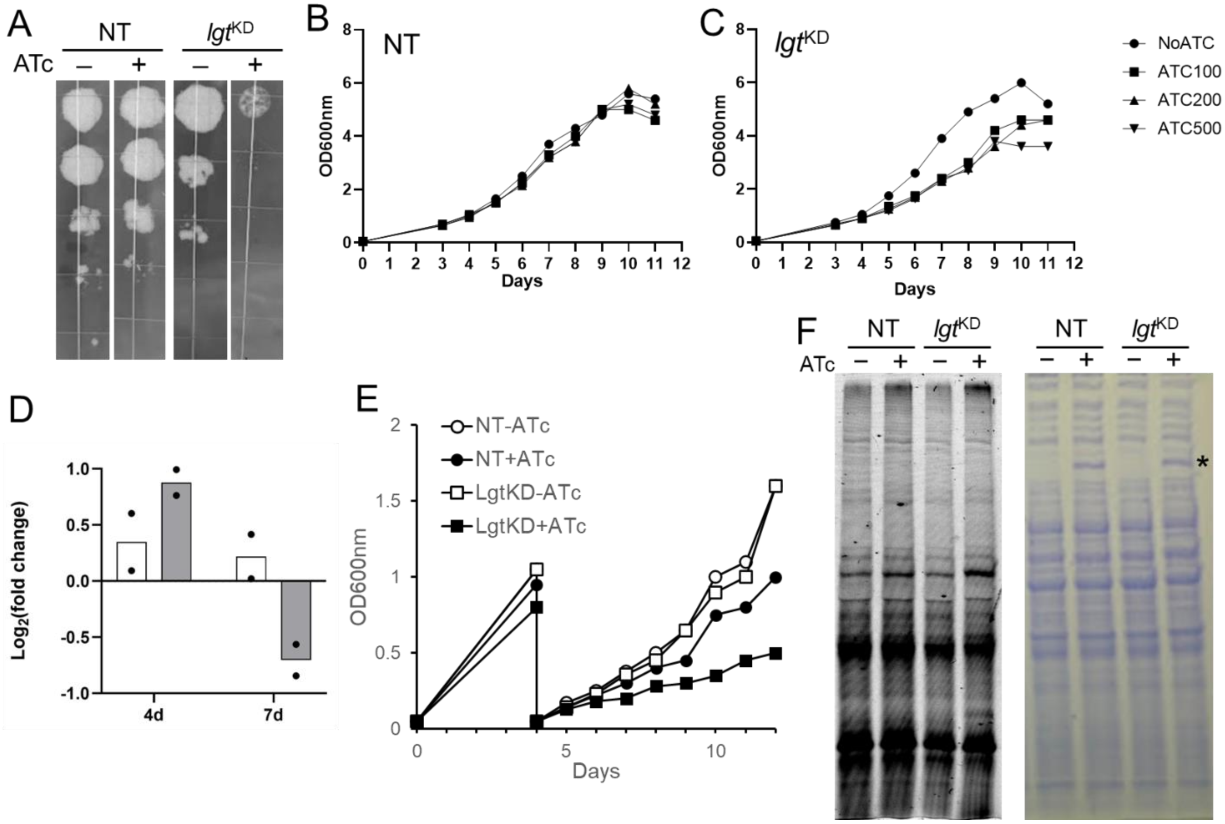


**Figure S5. L*gt* is essential but not highly vulnerable by CRISPRi knockdown and suppression does not affect protein labeling by BODIPY-C12 in *Mtb*.** A) *Mtb* encoding a scramble (NT, non-targeting) or *lgt*-targeting (*lgt*$^{KD}$) sgRNA was cultured to $OD_{600}$ ~ 1. A 10-fold dilution series was prepared starting from approximately 1000 CFU (5 µl culture diluted to $OD_{600}$ ~0.06) and spotted on Middlebrook 7H11 agar +/− anhydrotetracycline (ATc) at 100 ng/ml. Data were recorded after 15 d and are representative of n=2 independent experiments. Growth of B) NT and C) *lgt*$^{KD}$ strains without (NoATC) or with varying concentrations of ATc (100, 200, and 500 ng/mL) following inoculation at $OD_{600}$ 0.05. Data shown are representative of n=2 independent experiments. D) Total RNA was extracted from cultures in C), specifically *Mtb* NT (white bar) and *lgt*$^{KD}$ (grey bar) strains cultured −/+ ATc (0 or 100 ng/ml) for 4 or 7 days. Raw

Ct values for *lgt* (+ATc) were normalized to the housekeeping gene *sigA* and compared with −ATc control to calculate fold change. Data from n=2 biological replicates are shown. E) *Mtb* were cultured as in A), but subcultured after 4 days to enable incubation with ATc (100 ng/mL) while maintaining log-phase growth in control samples. F) Cultures from E) at day 12 were incubated with 5 µM BC12 for 20 h and compared by SDS-PAGE and fluorescence scanning. G) Coomassie blue staining did not reveal obvious proteome-wide perturbations upon addition of ATc; the induced band (*) is likely dCas9.

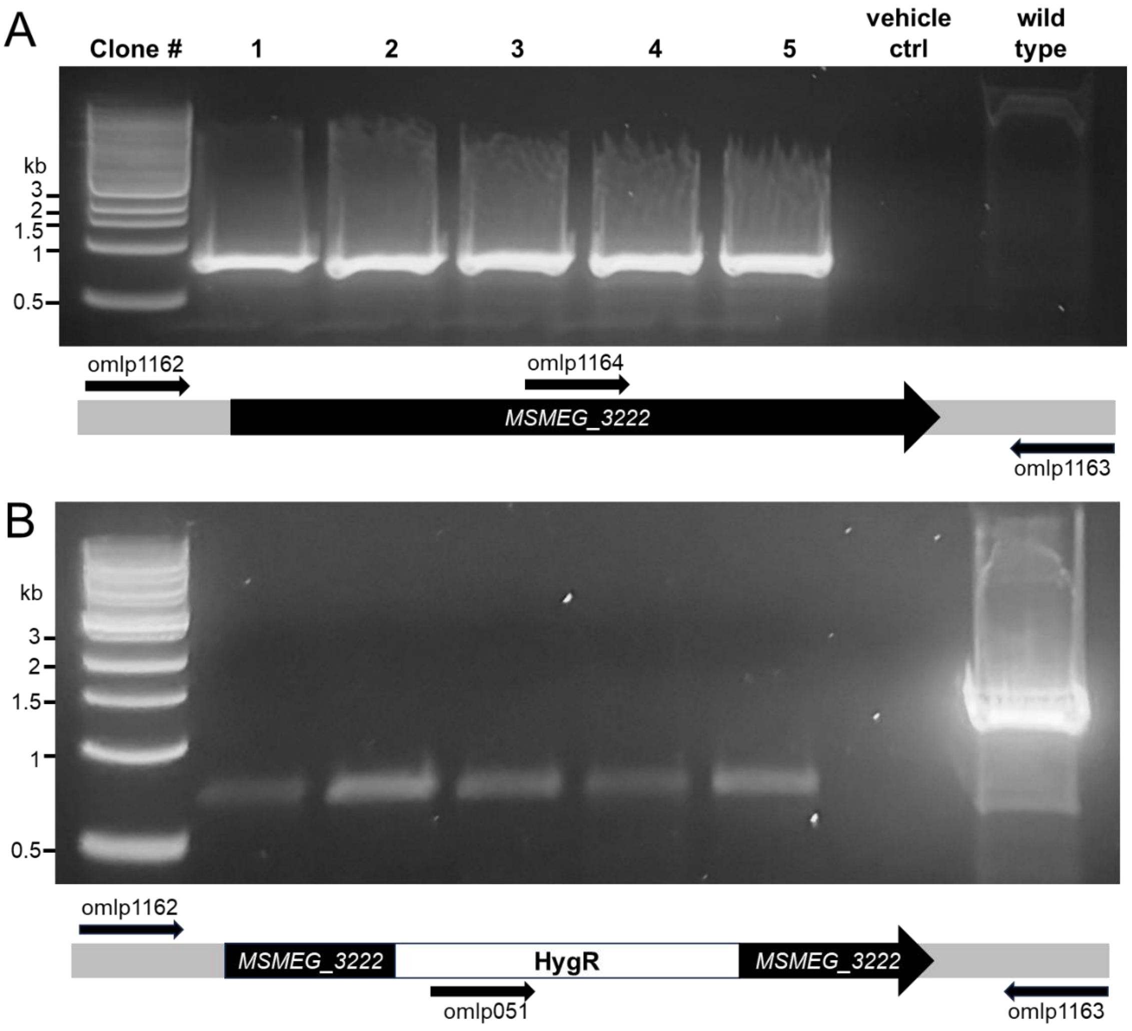


**Figure S6. Confirmation of Δ*MSMEG_3222* by PCR and sequencing.** Genomic DNA was isolated from five individual hygromycin-resistant clones as template for PCR with A) omlp051 and ompl1163 and B) omlp1164 and omlp1163 (**Table S2**). Either A) all five clones or B) only wild type yielded a product consistent with the expected size (A: 746 bp; B: 1430 bp). Water vehicle was used as a negative control. Clone 4 was selected for further characterization. Primers omp1162 and omlp1163 were used to generate a PCR product spanning the gene locus and the substitution was confirmed by sequencing.

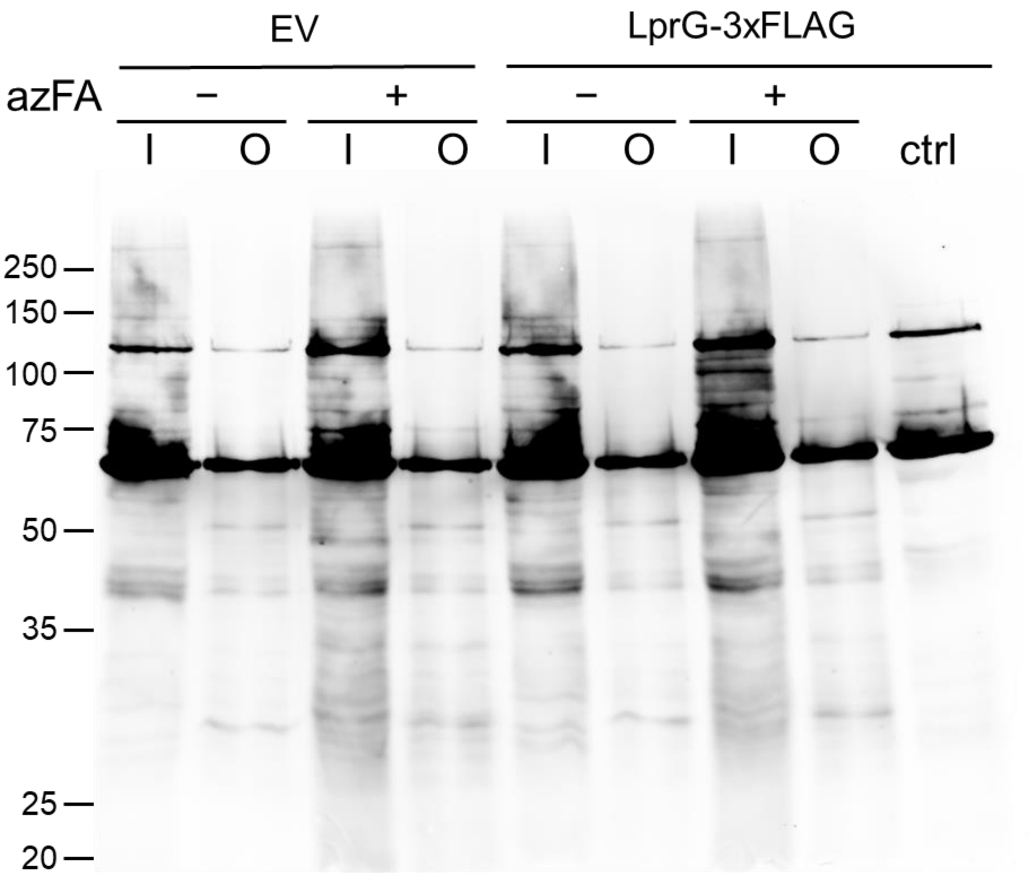


**Figure S7.** ***Msm*** **treatment with azFA followed by CuAAC to alk-TB and detection of biotin confirms azFA incorporation into proteins.** Biotinylated proteins in samples from Figure 4 were detected using strepavidin. Bands present in − azFA samples represent endogenously biotinylated proteins, as previously reported[1]. (I = input, O = output, ctrl = untreated total lysate from *Msm* LprG-3xFLAG)

### SUPPLEMENTAL METHODS

**Lipid extraction and analysis.** Following metabolic incorporation of BC12 into *Msm* as described in Methods (“Metabolic incorporation of fatty acids in *Msm*”), cells were washed and cultured for a further 2 h before harvesting by centrifugation. The cell pellet was resuspended in water and subjected to modified Bligh and Dyer extraction as follows: 1:2 chloroform:methanol (C:M) was added to a final ratio of 1:2:0.8 C:M:water (C:M:W) before storage at -20 °C until further processing. After samples were allowed to warm to 22 °C, chloroform and water were added with vortexing to obtain 1:1:0.8 C:M:W followed by centrifugation at 10,000 x *g* for 5 min. The lower phase was transferred to a fresh tube and allowed to dried overnight. The resulting film was resuspended in 2:1 C:M in 1/16th the original extraction volume.

A migration standard for TAG containing BC12 was generated from BC12-treated THP-1 cells as described[2,3]. Briefly, 3 x $10^6$ THP-1 monocytes were differentiated into macrophages using 100 µM PMA at a density of 0.6 million cells/ml for 24 h, followed by 2 days in media without PMA. BC12 was added to 1 µM and cells were cultured for an additional 24 h. Medium was removed and the monolayer was washed twice with PBS. Cells were then lysed in 0.5 mL PBS with 1% (v/v) TritonX-100. Four volumes of methanol:chloroform (2:1) was added to the lysate and vortexed for 30 seconds. One volume each of 50 mM citric acid, water, and chloroform were added with vortexing. After centrifugation at 10,000 x *g* for 10 min, the lower phase was isolated, dried and resuspended in 200 µL C:M (2:1).

For thin-layer chromatography, 5 µL of each sample was resolved on aluminum-backed TLC silica gel 60 plates (Supelco) in 100:14:0.8 C:M:W or 70:30:1

hexanes:diethyl ether:acetone and scanned for BODIPY fluorescence (Sapphire Bioimager, Azure Biosystems).

**Generation of a *Mtb lgt* CRISPRi knockdown strain.** A CRISPRi knockdown strain for *rv1614* was generated in *Mtb* as reported[4] (see **Tables S1-S3** for oligonucleotides used and plasmids and strains generated). Briefly, a PAM sequence was chosen based on its high predicted suppression strength (97%). Complementary oligonucleotides encoding the selected PAM sequence and flanking BsmBI enzyme sites were annealed and ligated into digested pIJR965. The resulting sequence-confirmed vector (*lgt*$^{KD}$) and pIJR965 as a non-targeting control (NT) were transformed into *Mtb* and individual clones selected on kanamycin for propagation in liquid medium. For initial cultures, 1 mL of a thawed frozen stock was inoculated into 5 mL culture medium and incubated for 3 d.

**Spot assay.** *Mtb lgt*$^{KD}$ and NT strains were subcultured to $OD_{600}$ ~ 1 and then diluted to $OD_{600}$ 0.06 in 1 ml of the same medium. The suspension was 10-fold serially diluted 5 times and 5 µl of each cell suspension was spotted on Middlebrook 7H10 agar with or without 100 ng/ml anhydrotetracycline (ATc). Data were recorded after 15 d incubation.

**RNA extraction and qRT-PCR.** For liquid cultures, *Mtb* was grown in 10 mL cultures overnight to $OD_{600}$ ~0.8-1. The cell pellet was resuspended in 1 mL of TRIzol LS (Invitrogen) and lysed with 0.1 mm zirconia beads by bead beating (Beadruptor, Omni International) for 30 s at 6 m/s followed by 5 min on ice for a total of 4 cycles. The lysate (~800 µl) was then thoroughly mixed with 200 µl chloroform:isoamyl alcohol. After centrifugation, the upper aqueous layer was transferred to a fresh tube and mixed with

360 µl ethanol. RNA was then purified using columns according to the manufacturer's instructions (Qiagen RNeasy kit) and DNA contamination removed (TURBO DNA-free kit, Invitrogen). DNA removal was confirmed by qPCR with *sigA* primers (**Table SX**). Purified RNA (~500-1000 ng) was then used to synthesize cDNA (Verso cDNA synthesis kit, Thermo Scientific). The cDNA was diluted 10-fold and 2 µL of the dilution was used for qRT-PCR analysis (Power SYBR Green PCR Master Mix, Applied Biosystems) on a LightCycler 480 Instrument (Roche). Ct values were calculated by LightCycler 480 Software (Roche) and normalized using *sigA* as a housekeeping gene. Primer efficiencies were verified by standard curves. Gene expression fold change was calculated using the ΔΔCt method[5].
